# Low-rate firing limit for neurons with axon, soma and dendrites driven by spatially distributed stochastic synapses

**DOI:** 10.1101/669705

**Authors:** Robert P. Gowers, Yulia Timofeeva, Magnus J. E. Richardson

**Affiliations:** Mathematics for Real-World Systems Centre for Doctoral Training, University of Warwick, Coventry, CV4 7AL, United Kingdom; Department of Computer Science, University of Warwick, Coventry, CV4 7AL, United Kingdom; Warwick Mathematics Institute, University of Warwick, Coventry, CV4 7AL, United Kingdom; Department of Clinical and Experimental Epilepsy, UCL Queen Square Institute of Neurology, University College London, London, WC1N 3BG, United Kingdom

## Abstract

Analytical forms for neuronal firing rates are important theoretical tools for the analysis of network states. Since the 1960s, the majority of approaches have treated neurons as being electrically compact and therefore isopotential. These approaches have yielded considerable insight into how single-cell properties affect network activity; however, many neuronal classes, such as cortical pyramidal cells, are electrically extended objects. Calculation of the complex flow of electrical activity driven by stochastic spatio-temporal synaptic input streams in these structures has presented a significant analytical challenge. Here we demonstrate that an extension of the level-crossing method of Rice, previously used for compact cells, provides a general framework for approximating the firing rate of neurons with spatial structure. Even for simple models, the analytical approximations derived demonstrate a surprising richness including: independence of the firing rate to the electrotonic length for certain models, but with a form distinct to the point-like leaky integrate-and-fire model; a non-monotonic dependence of the firing rate on the number of dendrites receiving synaptic drive; a significant effect of the axonal and somatic load on the firing rate; and the role that the trigger position on the axon for spike initiation has on firing properties. The approach necessitates only calculating first and second moments of the non-thresholded voltage and its rate of change in neuronal structures subject to spatio-temporal synaptic fluctuations. The combination of simplicity and generality promises a framework that can be built upon to incorporate increasing levels of biophysical detail and extend beyond the low-rate firing limit treated in this paper.

**Author summary:** Neurons are extended cells with multiple branching dendrites, a cell body and an axon. In an active neuronal network, neurons receive vast numbers of incoming synaptic pulses throughout their dendrites and cell body that each exhibit significant variability in amplitude and arrival time. The resulting synaptic input causes voltage fluctuations throughout their structure that evolve in space and time. The dynamics of how these signals are integrated and how they ultimately trigger outgoing spikes have been modelled extensively since the late 1960s. However, until relatively recently the majority of the mathematical formulae describing how fluctuating synaptic drive triggers action potentials have been applicable only for small neurons with the dendritic and axonal structure ignored. This has been largely due to the mathematical complexity of including the effects of spatially distributed synaptic input. Here we show that in a physiologically relevant, low-firing-rate regime, an approximate, level-crossing approach can be used to provide an estimate for the neuronal firing rate even when the dendrites and axons are included. We illustrate this approach using basic neuronal morphologies that capture the fundamentals of neuronal structure. Though the models are simple, these preliminary results show that it is possible to obtain useful formulae that capture the effects of spatially distributed synaptic drive. The generality of these results suggests they will provide a mathematical framework for future studies that might require the structure of neurons to be taken into account, such as the effect of electrical fields or multiple synaptic input streams that target distinct spatial domains of cortical pyramidal cells.

## Introduction

Due to their extended branching in both dendritic and axonal fields many classes of neurons are not electrically compact objects, in that the membrane voltage varies significantly throughout their spatial structure. A case in point are the principal, pyramidal cells of the cortex that feature a long apical dendritic trunk, oblique dendrites, apical tuft dendrites and a multitude of basal dendrites. Excitatory synapses are typically located throughout the dendritic arbour [1], while inhibitory synapses are clustered at specific regions depending on the presynaptic cell type [2]. These cells also differ morphologically not only between different layers, but also between cells in the same layer and class [3, 4]. Most cortical pyramidal cells *in vivo* fire rarely and irregularly due to the stochastic and balanced nature of the synaptic drive [5, 6]. Despite the apparent irregular firing of single neurons, computational processes are understood to be distributed across the population [7, 8] with the advantage that encoding information at a low firing rate can be energy efficient [9].

The arrival of excitatory and inhibitory synaptic pulses increases or decreases the postsynaptic voltage as well as increasing the conductance locally for a short time. Together with the spatio-temporal voltage fluctuations caused by the distributed synaptic bombardment typical of in vivo conditions, the increase in membrane conductance affects the integrative properties of the neuron, with reductions of the effective membrane time constant, electrotonic length constant and overall input resistance of neuronal substructures [10–12].

How different classes of neurons integrate stochastic synaptic input has been a subject of intense experimental [7, 13, 14] and theoretical [15–19] focus over the last 50 years. The majority of analytical approaches have approximated the cell as electrotonically compact and focussed on the combined effects of stochastic synaptic drive and intrinsic ion currents on the patterning of the outgoing spike train. Such models usually utilise an integrate-and-fire (IF) mechanism with some variations, and have been analysed using a Fokker-Planck approach [20–23] in the limit of fast synapses. However, this approach becomes unwieldy when synaptic filtering is included (though see [24–26]). One approximate analytical methodology, applicable to the low-firing-rate limit driven by filtered synapses is the level-crossing method of Rice [27]. In this approach, which has already been applied to compact neurons [28, 29], a system without post-spike reset is considered with the rate that the threshold is crossed from below treated as a proxy for the firing rate; the upcrossing rate and firing rate for a system with reset will be similar when the rate is low.

Due partly to the sparsity of the experimental data required for model constraint, but also because of the mathematical complexity involved, few analytical results regarding stochastic synaptic integration are available for neurons with dendritic structure, excepting the work of Tuckwell [30–32]. Nevertheless there is increasing interest in the integrative and firing response of spatial neuron models [33, 34], for neurons subject to and generating electric fields [35–37] or the effect of axonal load and position of the action-potential initiation region [34, 38–42]. Advances in optogenetics and multiple, parallel intracellular recordings have made experimental measurement and stimulation of *in vivo*-like input at arbitrary dendritic locations feasible [43–45]. This potential for model constraint suggests it is timely for a concerted effort to extend the analytical framework developed for compact models driven by stochastic synapses to neurons with dendrites, soma and axon in which the voltage fluctuates in both space and time.

Here we detail an analytical framework for approximating the firing rate of neurons with a spatially extended structure in a physiologically relevant low-rate regime [46–48]. To illustrate the approach we applied it to simple but exemplary neuronal geometries with increasing structural features - multiple dendrites, soma and axon - and investigated how various morphological parameters including the electrotonic length, axonal radius, number of dendrites and soma size affect the firing properties.

## Materials and Methods

### Derivation of the stochastic cable equation

The cable equation for the voltage *V*(*x, t*) in a dendrite of constant radius a and axial resistivity *r_a_* with leak and synaptic currents has the form

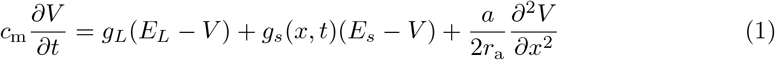

where *c*_m_, *g_L_* and *g_s_* are the membrane capacitance, leak conductance and synaptic conductance per unit area respectively, while *E_L_* and *E_s_* are the equilibrium potentials for the leak and synaptic currents. The synaptic conductance over a small area of dendrite, 2*πa*Δ_*x*_, at location *x* along the dendrite increases instantaneously by an amount *γ_s_* for each incident synaptic input and then decays exponentially with time constant *τ_s_* as the constituent channels close

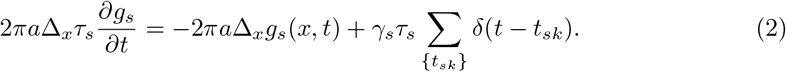

Here {*t_sk_*} denotes the set of synaptic arrival times at location *x*. Each synaptic pulse is assumed to arrive independently, where the number that arrive in a time window Δ_*t*_ is Poisson distributed with a mean *N_s_* given in terms of the dendritic section area, areal density of synapses *ϱ_s_*, and mean synaptic arrival rate *r_s_*

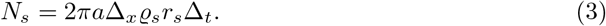

Note that for a Poisson process the variance will also be *N_s_*.

### Gaussian approximation for the fluctuating conductance

For a high synaptic-arrival rate we can approximate the Poissonian impulse train by a Gaussian random number with mean *N_s_*/Δ_*t*_ and standard deviation 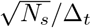 (this is an extension to spatio-temporal noise of the approach taken in [20]). Dividing Eq (2) by the unit of membrane area and introducing *ψ* as a zero-mean, unit-variance Gaussian random number allows us to write

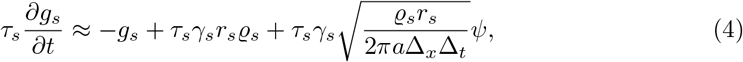

where the right-hand side should be interpreted as having been discretized over time, with a time step Δ_*t*_. We now define the space-time white-noise process 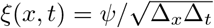 that has the properties

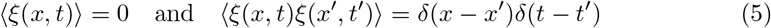

and also note that in the steady state 〈*g_s_*〉 = *τ_s_γ_s_r_s_ϱ_s_*. Returning to the cable equation, we split *g_s_* and *V* into mean and fluctuating components with *g_s_* = 〈*g_s_*〉 + *g_sF_* and *V* = 〈*V*〉 + *υ_F_*, giving the equation for the mean components as

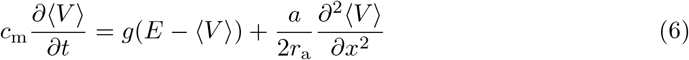

with *g = g_L_* + 〈*g_s_*〉 and *E* = (*g_L_E_L_* + 〈*g_s_*〉*E_s_*)/*g*. It is useful to introduce the time and space constants

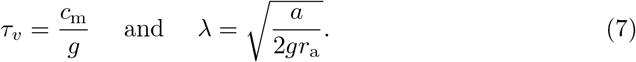

For the fluctuating component we assume that the product *g_sF_υ_F_* is small and obtain

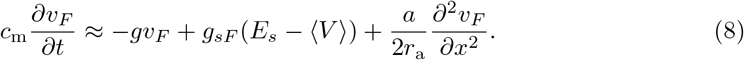

Rescaling synaptic variables *s = g_sF_*(*E_s_* − 〈*V*〉) and 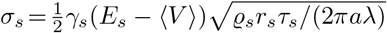 results in the following form for the synaptic equation

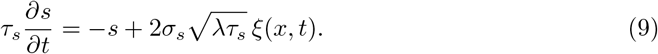

The deterministic voltage 〈*V*〉 is generally spatially varying. However, if the synaptic equilibrium potential *E_s_* is far from the effective resting voltage *E* and the fluctuating voltage remains close to *E* then it is reasonable to approximate the noise *σ_s_* as being spatially uniform with *E_s_* − 〈*V*〉 ≈ *E_s_* − *E*. This is applicable for mostly excitatory synaptic drive where *E_s_* ~ 0mV and *E* ~ −60mV. Letting *υ* = 〈*V*〉 − *E_L_ + υ_F_, μ = E − E_L_*, and substituting in *τ_υ_* and λ, we combine Eqs (6, 8) and (9) to obtain the stochastic cable equation used in the paper

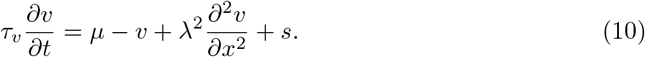

Here *μ* and *s* comprise the constant and fluctuating inputs to the dendrite. These subthreshold dynamics are supplemented by the standard integrate-and-fire threshold-reset mechanism at a trigger position *x*_th_; when the voltage at *x*_th_ exceeds a threshold *υ*_th_ the voltage in the entire structure is reset to voltage *υ*_re_. Under *in vivo* conditions the action-potential will back-propagate throughout the neuron with complex spatio-temporal dynamics [49–51]; however, here we are considering the low-rate case in which these transient post-spike dynamics will have dissipated before the next action potential is triggered.

### Boundary conditions

The morphologies explored in this paper are shown in Fig 1a and feature boundary conditions in which multiple dendrites and an axon meet at a soma. To account for these conditions we first define the axial current *I_a_* in a cable, writing it in terms of the input conductance of an infinite cable G_λ_ = 2*πa*λ*g*,

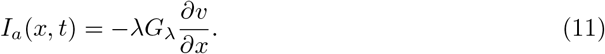

**Fig 1.**
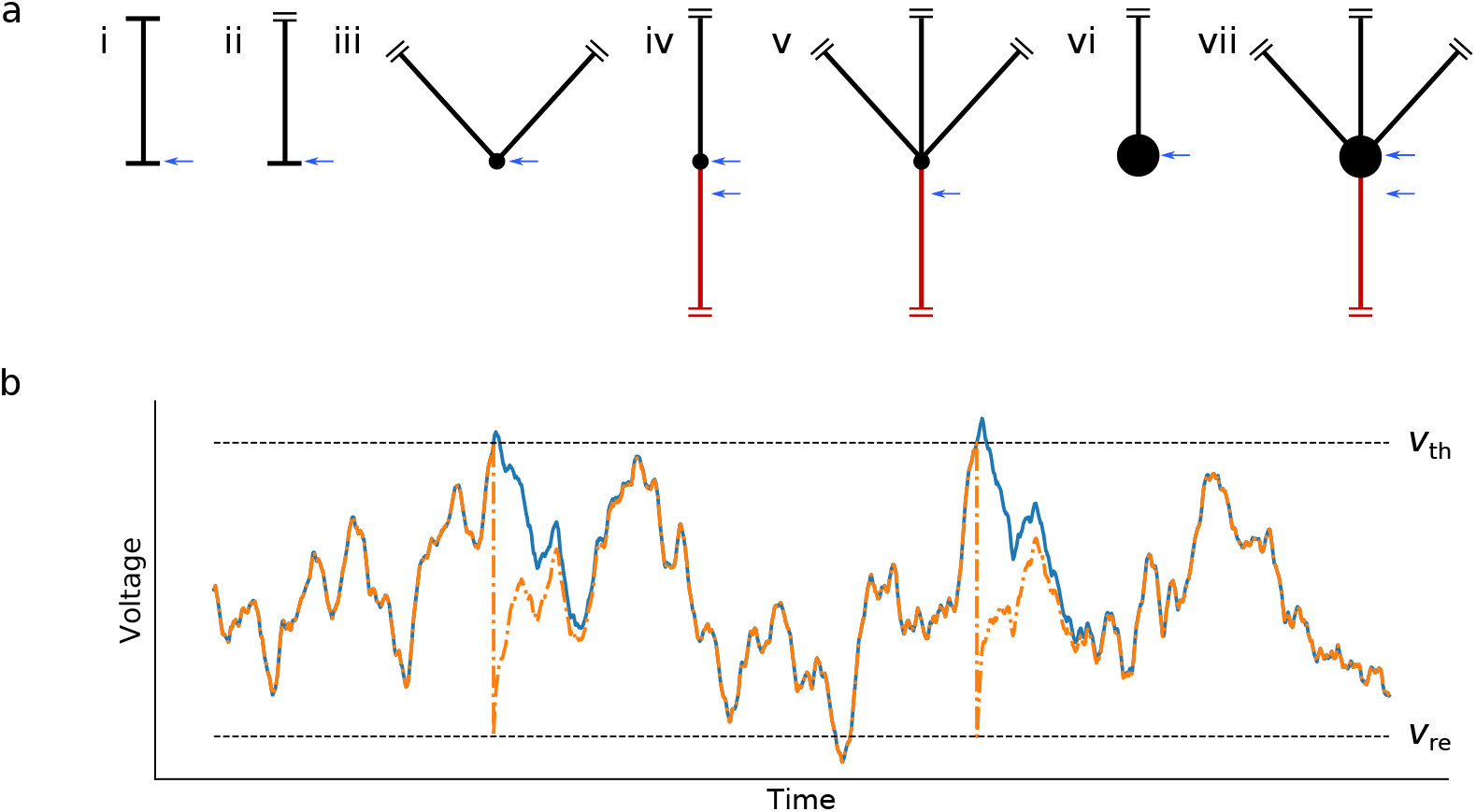
(a) The morphologies examined in this paper: (i) closed dendrite, (ii) semi-infinite one-dendrite model, (iii) two-dendrite model, (iv) dendrite and axon, (v) multiple dendrites and axon, (vi) dendrite and soma, (vii) multiple dendrites, soma and axon. Long black lines denote dendrites, red lines indicate the axon, while the blue arrows indicate the different spike trigger positions used. The symbols in the diagrams illustrate the following features: horizontal line - sealed end, two parallel lines - semi-infinite cable, small circle - nominal soma, and large circle - electrically significant soma. (b) Illustration of the upcrossing approximation. If the time between firing events is long compared to the relaxation time, the voltage without reset (solid blue line) will converge to the voltage of a threshold-reset process (orange dashed line) for the same realisation of stochastic drive. Under these conditions the upcrossing and firing-rates for the two processes are comparable.

For a sealed end at *x* = 0, represented by a horizontal line in Fig 1a, no axial current flows out of the cable giving the boundary condition

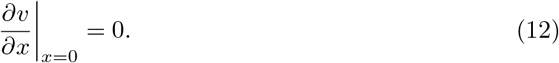

When the cable is unbounded and semi-infinite in extent, as shown by two small parallel lines in Fig 1a, we apply the condition that the potential must be finite at all positions,

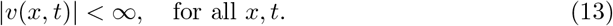

For other cases, multiple (*n*) neurites join at a nominal soma *x* = 0 which is treated as having zero conductance - these cases are shown by a small circle in Fig 1a. Under these conditions the voltage is continuous at the soma *υ*_1_ (0) = … = *υ_n_* (0) and axial current is conserved

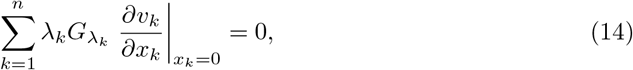

where *k* identifies the *k*th of the *n* neurites and *G*_λ*k*_ is its input conductance. Note that for each neurite the spatial variable *x_k_* increases away from the point of contact *x_k_* = 0. The addition of an axon changes this boundary condition by adding a cable of index *α* with length constant λ_*α*_ and conductance *G*_λ*α*_. Finally, when the soma at *x* = 0 is electrically significant (denoted by a large circle in Fig 1a), there is an additional leak and capacitive current at *x* = 0. This results in a current-conservation condition

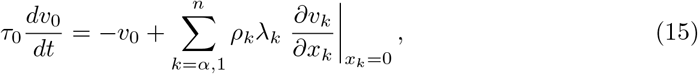

where the subscript 0 denotes somatic quantities and the neurite dominance factor *ρ_k_*, which is the conductance ratio between an electrotonic length of cable and the soma [52] is *ρ_k_* = *G*_λ*k*_/*G*_0_. As in the case for the nominal soma, the other condition is that the voltage is continuous.

### Numerical simulation

The cable equations for each neurite with a threshold-reset mechanism were numerically simulated by implementing the Euler-Maruyama method by custom-written code in the Julia language [53]. We discretized space and time into steps Δ_*x*_ and Δ_*t*_, with *υ* and *s* measured at half-integer spatial steps and the derivative *∂υ/∂_x_* at integer spatial steps. Hence, denoting *k* as the spatial index and *i* as the temporal index such that (*x, t*) = (*k*Δ_*x*_, *i*Δ_*t*_) 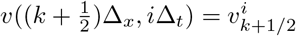 and 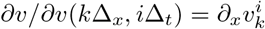. The numerical algorithm used to generate *υ* and *s* was therefore as follows

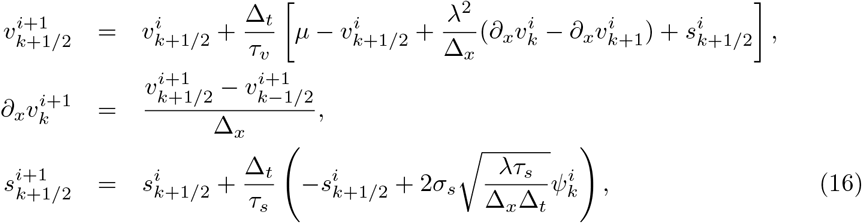

where *ψ* denotes a zero-mean, unit-variance Gaussian random number. The code used to generate the figures is provided in the supplementary information [54]. When the approximation of an infinite or semi-infinite neurite was required, the length L was chosen to be sufficiently large such that boundary effects were negligible (L = 1000*μ*m or greater). To ensure stability of the differential equation, for a spatial step of Δ_*x*_ = 20*μ*m, we used a time step of Δ_*t*_ = 0.02 ms. We verified that this step size was sufficiently small in comparison to the values of λ used by running simulations at smaller Δ_*x*_ and checking for convergence.

## Results

Before examining more complex spatial models with multiple dendrites, soma and axon, we first review the subthreshold properties of a single closed dendrite driven by fluctuating, filtered synaptic drive. We then illustrate how the upcrossing method can be applied to spatial models by interpreting the results for the closed dendrite as either a long dendrite with a nominal soma at one end or as two long dendrites meeting at a nominal soma. More complex neuronal geometries are then considered including those with multiple dendrites, axon and an electrically significant soma. The parameter ranges used are given in the Appendix in Table S1.

### Subthreshold properties of a closed dendrite

The dendrites considered here are driven by distributed, filtered synaptic drive. For reasons of analytical transparency, excitatory and inhibitory fluctuations are lumped into a single drive term *s*(*x, t*), though it is straightforward to generalise the synaptic fluctuations to two distinct processes. The fluctuating component of the synaptic drive obeys the following equation

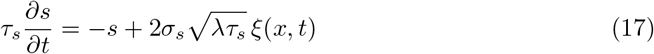

parametrized by a filter time constant *τ_s_*, amplitude *σ_s_* and driven by spatio-temporal Gaussian white noise *ξ*(*x, t*) (see Materials and Methods for links to underlying presynaptic rates and density, as well as the autocovariance of ξ(*x, t*)). Note that the fluctuating component of the synaptic drive *s*(*x, t*) is a temporally filtered but spatially white Gaussian process. The subthreshold voltage in the dendrite, driven by these synaptic fluctuations, will also be a fluctuating Gaussian process and obeys the following equation

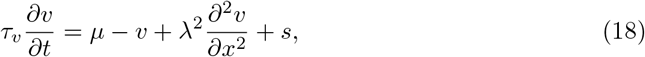

where the time constant *τ_υ_* and electrotonic length constant λ are reduced by the tonic conductance increase coming from the mean component of the synaptic drive (again, see Materials and Methods for derivation) and *μ* is the effective resting potential. For a closed dendrite of length *L*, shown in Fig 1a (i), there are two additional zero spatial-gradient conditions on *υ*(*x, t*) at *x*=0, *L*, Eq (12). With these definitions, it is straightforward to derive moments of the voltage using Green’s functions Eq (S21) into Eq (S14). The resulting second moments can then be more succinctly written by defining the function

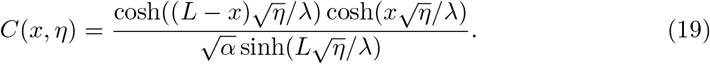

Hence in terms of this function *C*(*x, η*), the variance is

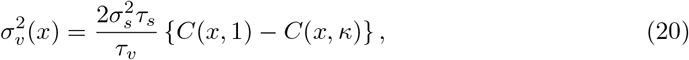

where *κ* = 1 + *τ_υ_/τ_s_*. Similarly from Eq (S15), the variance of 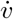 is

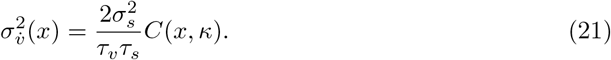

Note that the second term in the voltage-variance equation, Eq (20), and the variance of the voltage rate-of-change feature a second, shorter length constant 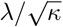 that is a function of the ratio of voltage to synaptic time constants. As expected, Fig 2a shows decreasing λ leading to a lower overall variance as well as a faster decay to the bulk properties from the boundaries. We also see from *κ* that the relative size of the time constants affects not just the magnitude of the variance but also its spatial profile. For higher 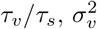 decreases at all positions and the profile decays faster to the bulk value as the second length constant decreases. By measuring 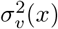 relative to the variance at the ends Fig 2b shows the latter effect, though this reduction in the effective length constant by increasing *τ_υ_/τ_s_* is not as significant as decreasing λ.

**Fig 2.**
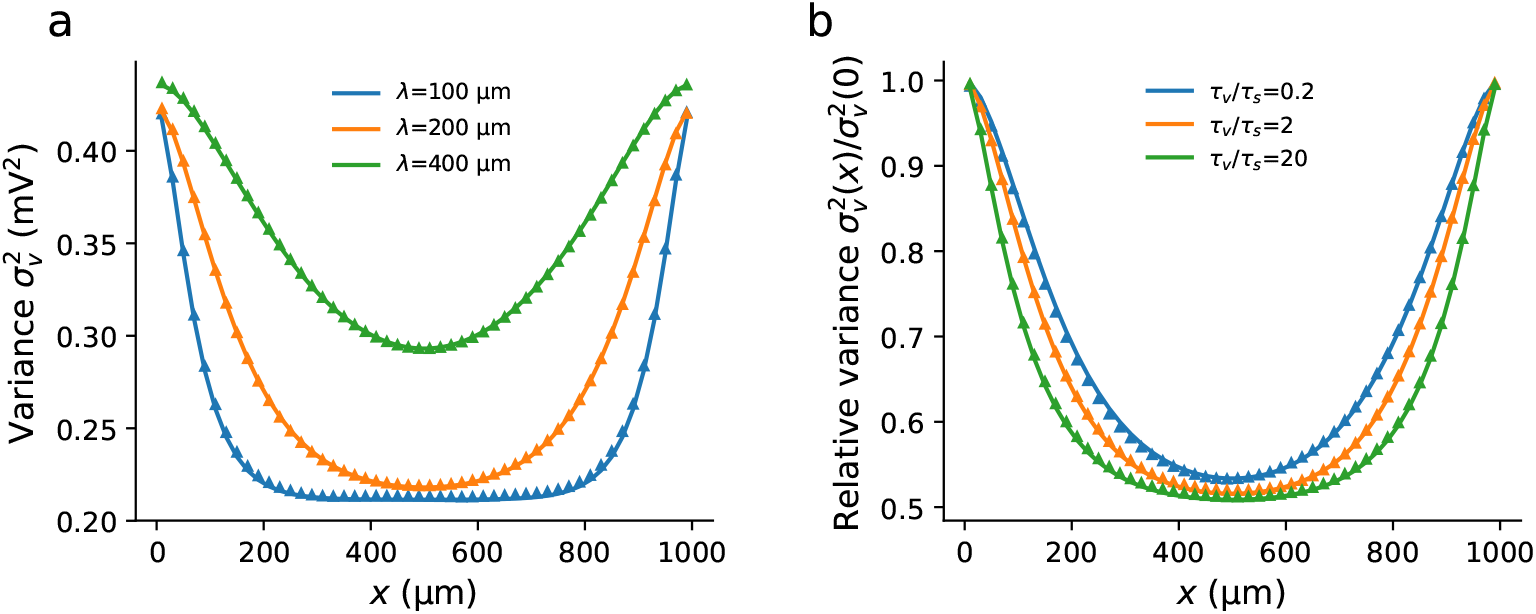
The variance profile in a sealed dendrite is a function of both the electrotonic length λ and ratio of synaptic to voltage time constants constant *τ_s_/τ_υ_*. (a) The variance near the sealed end is higher than in the bulk with the extent of the boundary effect decreasing with λ. (b) Normalising the variance in the cable such that the variance at the ends is unity, it can be seen that increasing *τ_υ_/τ_s_* decreases the effective length constant. For both plots the other parameters were *L* = 1000*μ*m, *τ_s_* = 5ms, *σ_s_* = 1mV. (a) *τ_υ_* = 10ms, (b) λ = 200*μ*m.

Note that for the cases where λ/*L* ≪ 1, which is physiologically relevant for the high-conductance state, the influence of the boundary at *L* is negligible at *x* = 0 and at the midpoint there is little influence from either boundary. With this in mind, the morphologies treated in this paper comprise neurites that are treated as semi-infinite in length.

### Firing rate approximated by the upcrossing rate

Full analytical solution of the partial differential Eqs (17, 18) when coupled to the integrate-and-fire mechanism does not appear trivial, even for the simple closed dendrite model. However, a level-crossing approach developed by Rice [27] and exploited in many other areas of physics and engineering, such as wireless communication channels [55], sea waves [56], superfluids [57] and grown-surface roughness [58] has previously been applied successfully to compact neuron models [28,29] and can be extended to spatial models. The method provides an approximation for the mean first-passage time for any Gaussian process in which the mean 〈*υ*〉, standard deviation *σ_υ_*, and rate-of-change standard deviation *σ_υ_* are calculable. The upcrossing rate is the frequency at which the trajectory of *υ* without a threshold-reset mechanism crosses *υ*_th_ from below (i.e. with 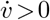).

Example voltage-time traces for the model with and without threshold are compared in Fig 1b. This approach provides a good approximation to the rate with reset when the firing events are rare and fluctuation driven, making it applicable to the physiologicallow-rate firing regime. The upcrossing approximation to the firing rate is given by

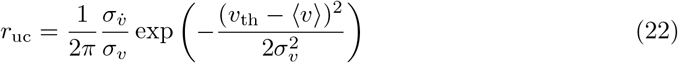

where the statistical measures of the voltage are those at the trigger point *x*_th_. Note that because of the requirement that 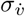 exists the upcrossing method cannot be applied to neurons driven by temporal white noise. However, it works well for coloured-noise drive, which is not directly tractable using standard Fokker-Planck approaches even for point-neuron models. The moments required for the upcrossing Eq (22) can be found using the Green’s functions of the corresponding set of cable equations for a particular morphology and, since we only need the moments at *x*_th_, we only need the Green’s function for the neurite that contains the trigger position (see Appendix for details). We now illustrate this using two interpretations of the closed dendrite model, the one-dendrite model which focuses on the behaviour at a sealed end - Fig 1a (ii) - and the two-dendrite model which focuses on the bulk - Fig 1a (iii).

### One-dendrite and two-dendrite models

The method is first applied to a neuron with a single long dendrite and nominal soma (the trigger point *x* = 0 = *x*_th_) and axon, both of negligible conductance so that the end can be considered sealed. This corresponds to the *x*<*L*/2 half of the closed cable model considered above, in the limit that *L*/λ→∞. The second moments have already been calculated for the general case (Eqs 20, 21) so for *x*_th_=0 we have

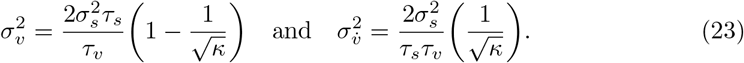

Substitution of these second moments into Eq (22) yields the upcrossing approximation to the firing rate for this geometry.

A second interpretation of the closed dendrite model is to place the trigger position in the middle *x*_th_=*L*/2 and then, in the limit *L*/λ→∞ consider the halves as two dendrites with statistically identical properties radiating from a nominal soma and axon, again both with negligible conductance. Taking these limits of the closed-dendrite Eqs (20, 21) for this case generates second moments that happen to be exactly half that of the one-dendrite case

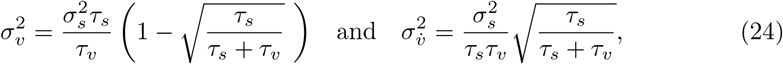

where here we have written the functional dependence of *κ* on *τ_υ_* and *τ_s_* explicitly. Given that the voltage at *x*_th_ is affected by activity occurring within distances a few λ down attached dendrites (see Fig 2) it might reasonably be expected that the statistical quantities and therefore the firing rate at *x*_th_ would be dependent on the electronic length quantity λ. However, for both the one and two-dendrite models considered above it is clear that there is no λ dependence for the second moments. Though this is unavoidable on dimensional grounds, because in either case no other quantities carry units of length once the limit *L*/λ→0 has been taken, the result is nevertheless a curious one.

The upcrossing and firing rates as a function of *μ* for the two models are compared in Fig 3, with the deterministic firing rate also shown (this is equivalent to the deterministic rate of the leaky integrate-and-fire model). Note that we keep *τ_υ_* and λ constant across the range of *μ* since these parameters would change little across the range of mean synaptic drive we investigate and it allows us to isolate the dependence of the firing rate on just one parameter. The upcrossing rate provides a good approximation to the full firing rate at low rates in the < 5Hz range. In this way the upcrossing rate for spatio-temporal models provides a similar approximation to the firing rate as did the Arrhenius form derived by Brunel and Hakim [20] for the white-noise driven point-like leaky integrate-and-fire model.

**Fig 3.**
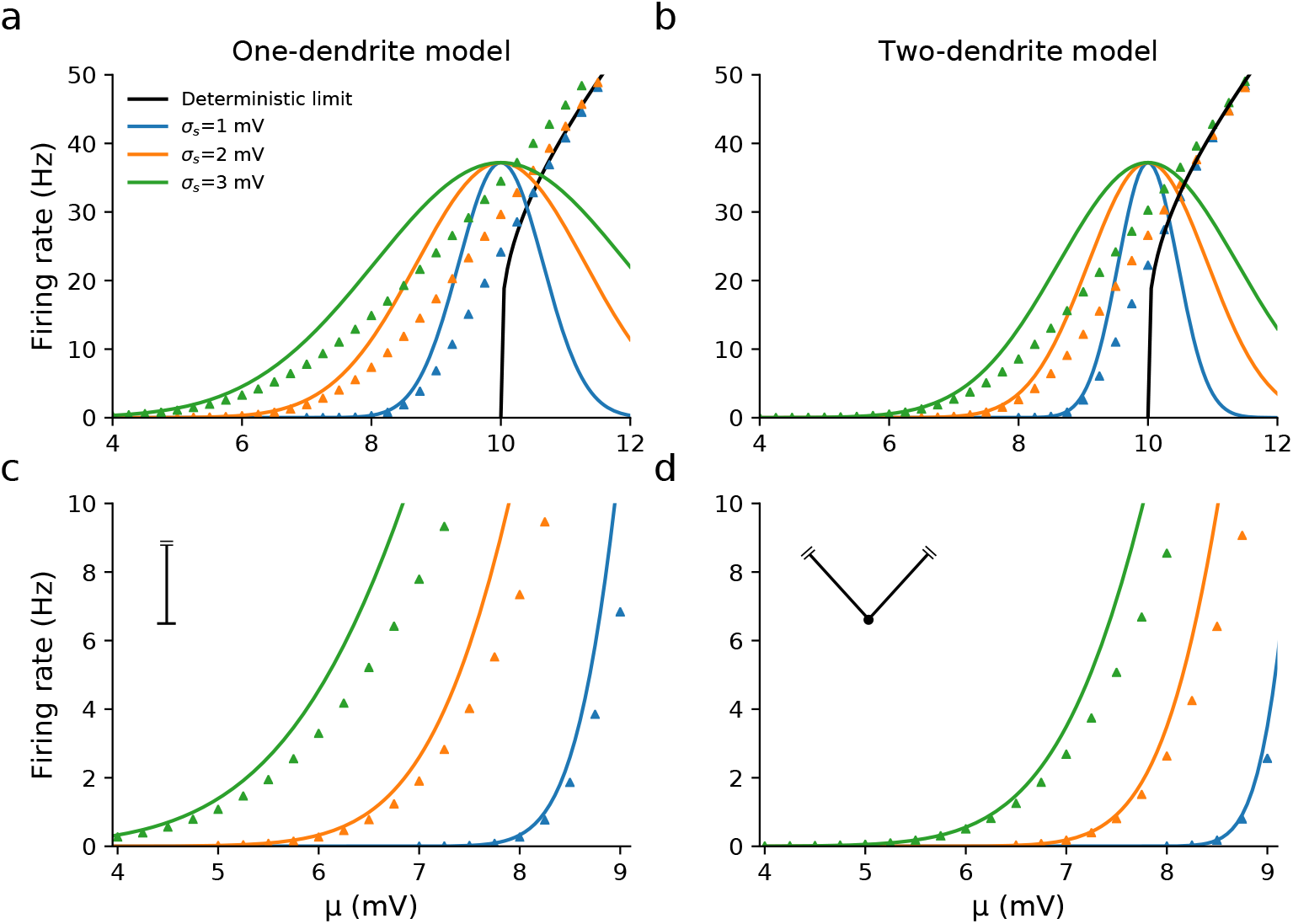
The firing rates of the one and two-dendrite models driven by spatially distributed, filtered stochastic synaptic drive for three fixed values of the fluctuation strength *σ_s_*. While the firing rates of the one and two-dendrite models are similar in the suprathreshold regime (panels a and b, *μ* > 10mV), the one-dendrite model has a higher firing rate in the subthreshold regime due to the variance being twice that of the two-dendrite model for the same value of *σ_s_*. The upcrossing approximation is accurate when (*υ*_th_ − *μ*)/*σ_υ_* ≫ 1 (panels c and d). The other parameters used were λ = 200*μ*m, *τ_υ_* = 10 ms, *τ_s_* = 5ms, *υ*_th_ = 10mV, *υ*_re_ = 0mV.

Compared with the one-dendrite model, we see from Figs 3b and 3d that the firing rate for the two-dendrite model is significantly lower in the subthreshold regime but converges to the same value when *μ* goes above threshold. This illustrates that even simple differences in morphology affect stochastic and deterministic firing very differently. In addition Fig 4a shows that the firing rate is unaffected by the value of λ chosen, confirming by simulation the λ-independence of the firing rate. Furthermore when we choose the same value of *σ_υ_* for the one and two-dendrite models, then both the upcrossing rate and the simulated firing rates are the same, as seen in Fig 4b.

**Fig 4.**
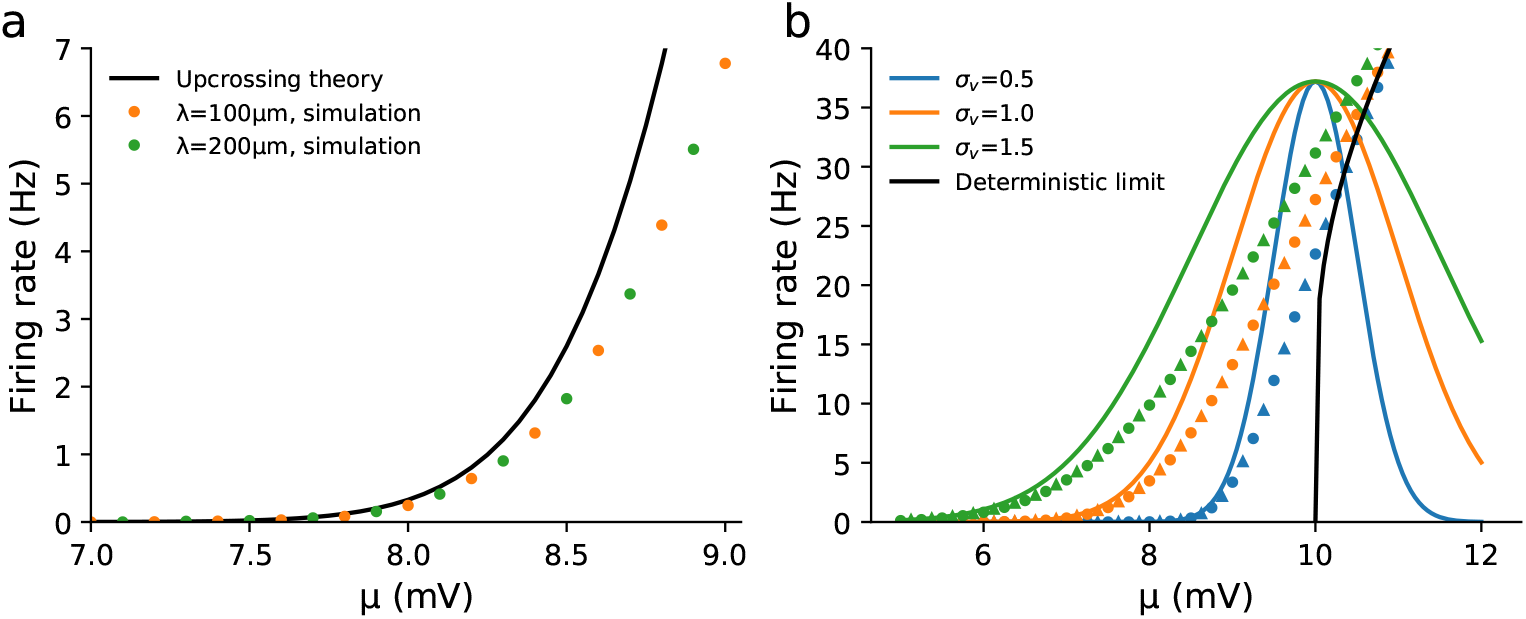
Independence of the firing rate on the electrotonic length λ for the one-dendrite model, and between the one and two-dendrite models for the same voltage variance. (a) The firing rate of the one-dendrite model with two different λ show it to be independent from λ, *σ_s_* = 1mV. (b) If the synaptic fluctuation strength *σ_s_* is adjusted such that the one and two-dendrite models have the same voltage variance 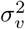 at the threshold position, then their upcrossing rates are identical. Simulations (circles and triangles for one and two-dendrite models respectively) suggest that the full firing rates are also independent of geometry in this case. The other parameters used were *τ_υ_* = 10ms, *τ_s_* = 5ms, *υ*_th_ = 10mV and *υ*_re_ = 0mV.

However, despite the independence of λ, the firing-rate profile for this toy model is distinct to that for the point-like leaky integrate-and-fire model, for which the second moments are 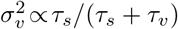 and 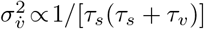 [29]. This indicates that spatial structure by itself decreases the variance while increases derivative variance by a factor 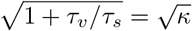. The moments also differ in their dependence on *τ_υ_* and *τ_s_* from two-compartmental models [59].

### Dendrite and axon

Next, we consider a dendrite connected to an axon at *x*_1_ =0 = *x_α_*, as shown in Fig 1a (iv), where dendritic and axonal quantities are denoted by subscripts 1 and *α*, respectively. This differs from the previous two-dendrite model as the axon receives no synaptic drive, so *μ_α_* = 0 and *s_α_* = 0. Furthermore, intrinsic membrane properties of the axon (*τ_α_*, λ_*α*_) differ from the dendrite due to the smaller axonal radius and lack of synapse-induced increased membrane conductance [11,12]. Since *μ_α_* = 0 we omit the subscript on the mean dendritic drive, *μ*_1_ = *μ*. Taking the reasonable assumptions that the per area leakage capacitance and leak conductance are the same in the axon as the soma, we can calculate *τ_α_* in terms of *τ*_1_ given the mean level of synaptic drive (see Eqs (S39, S41)). Unlike the closed-dendrite models, the mean is no longer homogeneous in space due to the lack of synaptic drive in the axon. Defining 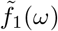 as the input admittance of the dendrite relative to the whole neuron,

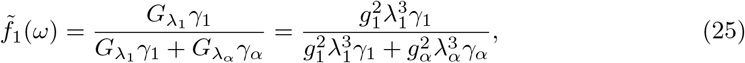

where 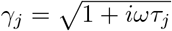, we can show that the mean in the axon is given by (see Eq (S12))

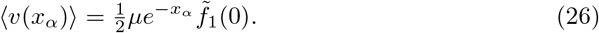

It is important to note that, unlike in the one and two-dendrite models, Eq (26) implies that it is now possible for the neuron to still be in the subthreshold firing regime when *μ* > *υ*_th_. In general, the second moments do not have a closed-form solution but can be expressed in terms of the angular frequency ω. It can be shown that the integrand for 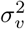 and 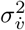 is proportional to 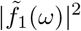, Eq (S38).

First we set the action-potential trigger position at *x*_th_ = 0 and evaluated the effect of the axon by comparing the firing rate for the model with an axon, *r*_axon_, to the firing rate of the one-dendrite model with a sealed end (*∂υ/∂x*=0) at *x*=0, *r*_sealed_ (effectively an axon with zero conductance load). We also kept the noise strength *σ_s_* rather than the voltage standard deviation *σ_υ_* fixed as we wished to see how the axon changes the variance of fluctuations at the trigger position. The relative firing rate was defined as *r*_axon_/*r*_sealed_. The ratio of the axonal to dendritic radius *a_α_*/*a*_1_ was varied and the relative firing rate calculated, with *a_α_*/*a*_1_=0 being equivalent to no axon present. As illustrated in Fig 5a, the addition of a very low conductance or relatively thin axon significantly reduces the firing rate. This effect arises because the magnitude of 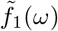 decreases at all frequencies for a larger radius ratio, which can be understood by recalling that 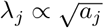, Eq (7).

**Fig 5.**
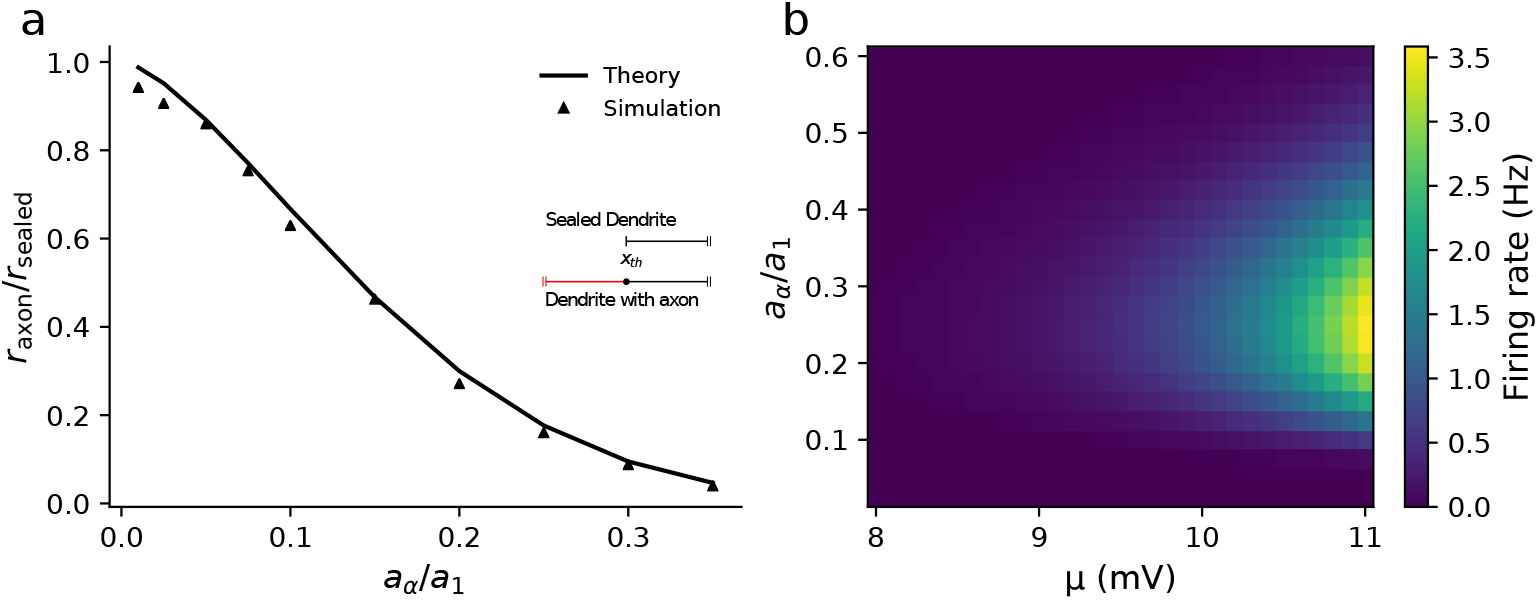
Addition of an axon significantly affects the firing rate even for small axonal conductance loads. (a) The axon increases the input conductance of the neuron, thereby lowering the firing rate for *x*_th_=0, *μ*=5mV. (b) When *x*_th_>0 (here *x*_th_=30μm) the firing rate varies non-monotonically with the axonal radius and peaks at a physiologically reasonable value of the ratio of axon/dendrite radii for a range of synaptic parameters. The parameters used were λ_1_=200*μ*m, *τ_s_* =5ms, *τ*_1_=10ms, *τ_α_*=11.7ms (calculated from Eq (S39)), *σ_s_*=3mV, *υ*_th_ =10mV and *υ*_re_=0mV.

For cortical pyramidal cells, action potentials are typically triggered around *x*_th_ = 30*μ*m down the axon in the axon initial segment [60–62]. It is straightforward to investigate the effect of moving the trigger position down the axon using the upcrossing approach. Interestingly, when *x*_th_ > 0, a non-monotonic relationship between the firing rate and radius ratio *a_α_*/*a*_1_ became apparent (see Fig 5b), with the peak ratio of ~ 0.25 being similar to that between the axonal initial segment and apical dendrite diameter in pyramidal cells [41,63]. This is caused by a non-monotonic dependence of both 〈*υ*〉 and 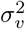 on *a_α_/a*_1_ for *x*_th_>0 with each peaking at intermediate values. Intuitively, this can be understood from the definition of λ_*α*_, which increases as 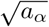. Thus the decay length of voltage fluctuations that enter the axon from the dendrite increases, increasing both 〈*υ*〉 and 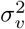 at *x*_th_. On the other hand, a larger λ_*α*_ increases the input conductance of the neuron, which, conversely, decreases 〈*υ*〉 and 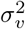. For smaller λ_*α*_ the decay length effect is more significant, whereas for larger λ_*α*_ the increase in input conductance plays a larger role.

### Multiple dendrites and axon

We now consider a case with multiple dendrites and an axon radiating from a nominal soma (Fig 1a (v)). The dendrites are labelled 1, 2, …, *n* with the axon labelled α as before. The dendrites have identical properties with independent and equally distributed synaptic drive. As in the previous case with the dendrite and axon, we keptthe synaptic strength *σ_s_* fixed as we changed the number of dendrites. An immediate consequence of multiple dendrites is that, since *μ* > 0 the mean voltage in the axon increases as more dendrites are added, with each contribution summing linearly,

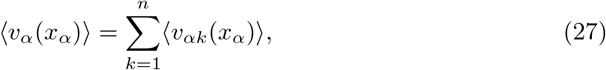

where 〈*υ*_*αk*_ (*x_α_*)〉 is the contribution to the axonal voltage mean from dendrite *k*. Introducing the relative input admittance of a single dendrite 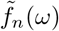

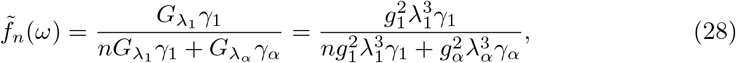

it can be shown that when all dendrites have identical mean input drive μ, the mean in the axon is given by (see Eqs (S12, S23))

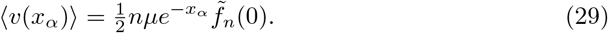

Thus we can see that as *n* increases the mean increases towards the constant value of 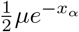. However, this is not the case for the fluctuating component: despite more sources of fluctuating synaptic input both 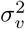 and 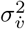 in the axon decrease as 1/*n* for a large number of dendrites. We can see this by noting that for large *n*, 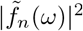 and hence the variance contribution from each dendrite scales as 1/*n*^2^. Therefore for *n* total dendrites, the total variance at *x*_th_ in the axon will scale as 1/*n* for large *n*. This reduction in axonal variance with additional dendrites is a generalisation of the reduction in variance we saw between the one and two-dendrite models earlier in Eqs (23, 24).

When it is the fluctuations that contribute significantly for firing (i.e. smaller *μ* or λ_*α*_) then a reduction in variance from adding more dendrites will decrease the firing rate; however, when the mean is more significant (larger *μ* or λ_*α*_) then the firing rate will increase as the number of dendrites increases. An example of the former case is shown in Fig 6a for λ_*α*_ = 100*μ*m, while an example of the latter is seen in Fig 6b for λ_*α*_ = 150*μ*m. The transition between these regimes can be seen in Fig 6c, which shows how the value of n that maximises the firing rate, *n*_max_, increases with *μ* and *a_α_*/*a*_1_. Physiologically, the reduction in variance is not simply the fact that adding dendrites increases cell size and thus input conductance, but that the relative conductance of each input dendrite to the total conductance decreases. Given that the total input conductance for *n* dendrites and an axon is

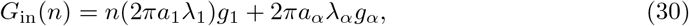

we can test this idea by scaling λ_1_, *a*_1_ with *n* (i.e. making the dendrites thinner) to keep the total input conductance the same as the single dendrite case, *G*_in_(*n* = 1). This gives the simple relationship λ_1_(*n*)=λ_1_(*n* = 1)/*n*^1/3^, with which the segment factor is

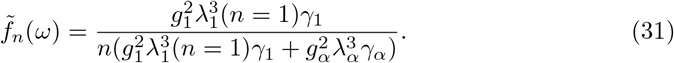

**Fig 6.**
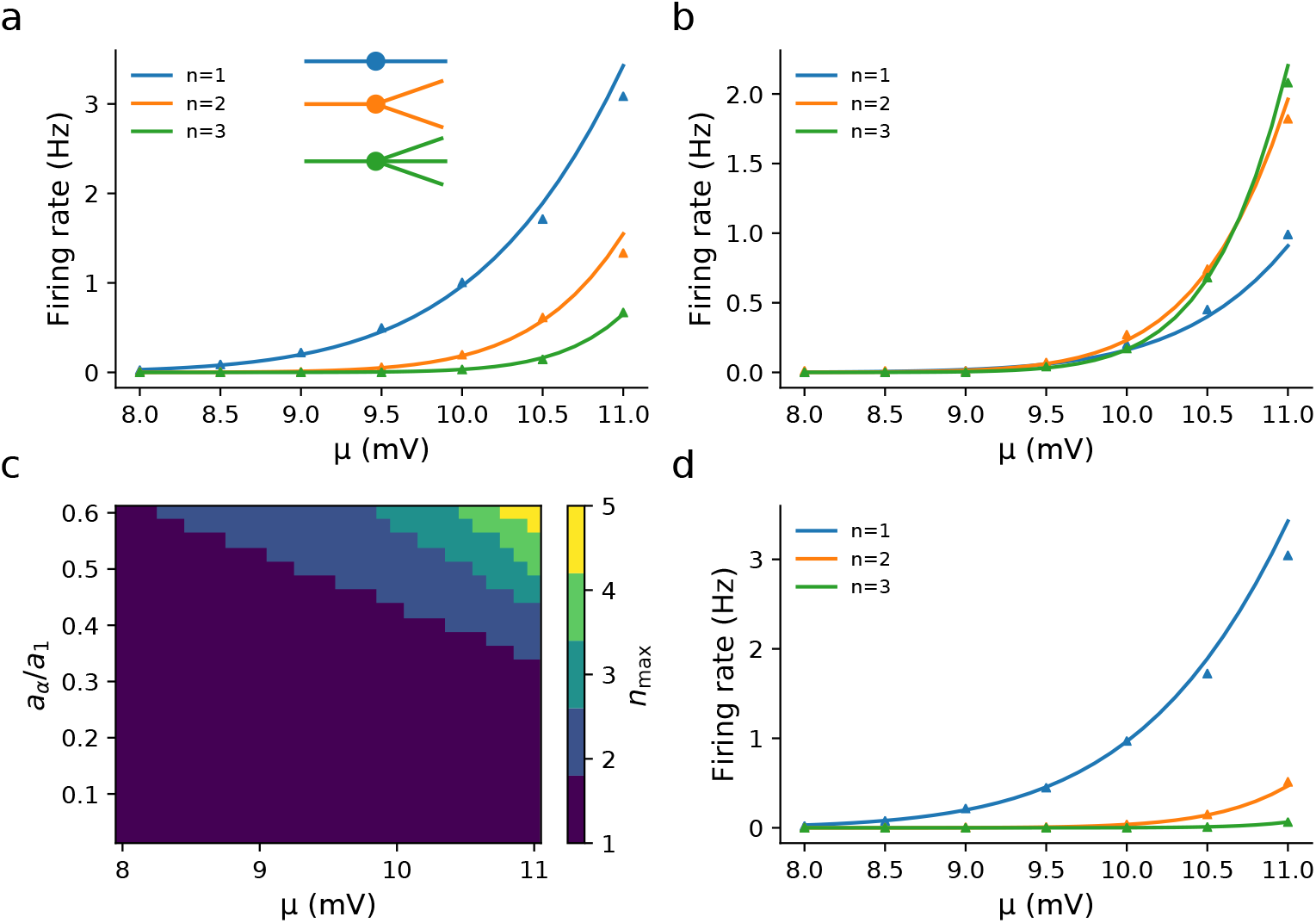
Increasing the number of synaptically driven dendrites can decrease the firing rate when the axon, of radius *a_α_* is much thinner than the dendrite, radius *a*_1_. The length constant for each neurite is proportional to the square root of the radius 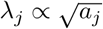. (a) λ_*α*_ = 100*μ*m, (b) λ_*α*_ = 150*μ*m, (c) The number *n*_max_ of dendrites that maximises firing increases with higher ratios of axon-to-dendrite radii *a_α_*/*a*_1_ and *μ*, (d) λ_*α*_ = 100*μ*m, λ_1_ = 200*μ*m for *n* = 1 and λ_1_ is rescaled for larger *n* to keep the input conductance equal to the *n* = 1 case. Other parameters: λ_1_ = 200*μ*m (a-c), *τ*_1_=10ms, *τ_α_*=11.7ms (calculated from Eq (S39)), *σ_s_* = 3mV, *υ*_th_=10mV and *x*_th_=30*μ*m down the axon.

Since the integrands for the second moments are proportional to 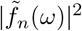 (see Eq (S38)), this shows that second moments and hence the firing rate for fixed λα still decrease with *n* (see Fig 6d).

### Dendrites, soma and axon

We now consider the case illustrated in Fig 1a (vii), where the electrical properties of the soma are non-negligible with its lumped capacitance and conductance providing an additional complex impedance at the point where the axon and dendrites meet. This has the somatic boundary condition we gave earlier in Eq (15) and we recall that the subscript 0 denotes somatic quantities. For simplicity, and as neither section receives synaptic drive in our model, we will let the somatic time constant be the same as the axonal time constant, so *τ*_0_ = *τ_α_*. Note that somatic drive can be straightforwardly added in this framework, as the variance contribution from the resultant fluctuations would add linearly: this would not qualitatively change the nature of the results we present here that focus on the effects of somatic filtering on transfer of dendritic stimulation to the trigger point in the axon. Note also that as *ρ*_1_ → ∞, the model without somatic drive converges to the dendrite and axon model with a nominal soma, allowing a clearer comparison between the two models.

For an electrically significant soma the integrand for the variance has the same form as before, Eq (S38), but 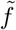 now depends on the neurite dominance factor *ρ*,

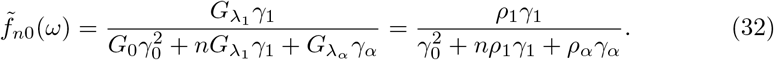

Thus for large *n* we should expect the variance in the axon to scale as 1/*n* as before, but for smaller n the somatic impedance 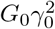 gives some key differences. We repeated the simulations for the axon-dendrite model (Fig 6), first with a single dendrite and an electrically significant soma by varying *ρ*_1_, noting that with known λ_1_ and λ_*α*_, this also determines *ρ_α_*, Eqs (S45, S46). Since the soma adds a conductance load *G*_0_ to the cell the overall input resistance decreases. From Eq (32), we see that this will reduce 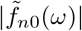, for any number of dendrites which will lower both the mean and the variance. Fig 7a shows that the effect of a larger soma (lower *ρ*_1_) lowers the firing rate.

**Fig 7.**
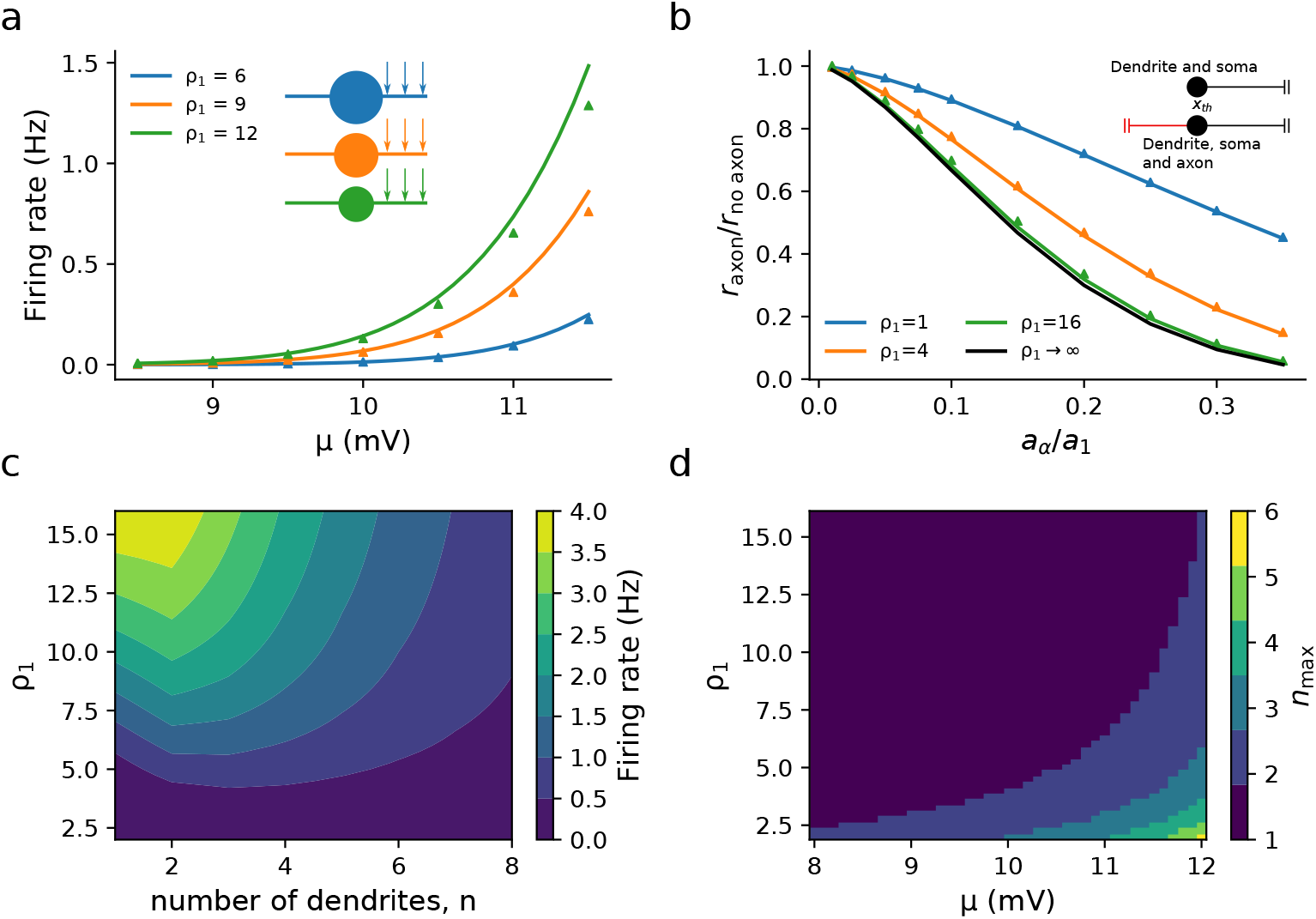
Effect of somatic impedance between a synaptically driven dendrite and axon with trigger point *x*_th_=30*μ*m for (a), (c) and (d) and *x*_th_=0*μ*m for (b). (a) The soma is characterised by the dendritic dominance factor *ρ*_1_ (see main text) with large *ρ*_1_ corresponding to a small somatic conductance. (b) A larger soma masks the effect of the axonal load on the firing rate, although this masking is negligible for smaller somata *ρ*_1_ = 16. Fig 2a (black line) result is plotted for comparison. (c) Larger somata also reduce the firing rate in the case of n dendrites (*μ* = 12mV). (d) With larger *μ*, smaller *ρ*_1_ (a larger soma) increases the number of dendrites for which the firing rate is maximal, *n*_max_. Other parameters: *τ*_1_ =10ms, *τ_α_*=11.7ms, λ_1_ =200*μ*m, λ_*α*_ =100*μ*m (calculated from Eq (S39)), *σ_s_*=3mV, *υ*_th_=10mV, *ρ_α_* calculated from Eqs (S45) or (S46)

Next, we investigated the effect an electrically significant soma has on axonal load, as seen previously for a nominal soma in Fig 5a. Like with the nominal soma case before, we calculated the firing rate at *x*_th_ = 0 with an axon and electrically significant soma, *r*_axon_, and the firing rate of a dendrite with the same size soma without an axon, *r*_no axon_ (Fig 1a (vi)). For each somatic size, we adjusted *σ_s_* so that the firing rate for a negligible axon, *a_α_*/*a*_1_ =0, the firing rate was fixed at 1Hz. This was done to account for the soma’s effect on the firing rate we observed earlier and we are thus solely focusing on the effect of the axonal admittance load. As we increase *a_α_*/*a*_1_ =0 Fig 7b shows that *r*_axon_/*r*_no axon_ decreases more rapidly with increasing *a_α_/a*_1_ for larger *ρ*_1_ (smaller soma). This means that, in comparison to Fig 5a, the axonal load had a lower relative effect on the firing rate in the presence of a soma. This is in line with what we should expect by looking at 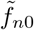; lower *ρ*_1_ increases the relative magnitude of 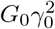 in the denominator of 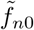 as compared with the axonal admittance term of *G*_λ*α*_ γ_*α*_.

Finally, we looked at how an electrically significant soma affects the dependence of the firing rate on the number of dendrites. By varying *ρ*_1_ and the number of dendrites *n*, Fig 7c shows that the non-monotonic dependence of the firing rate on dendritic number n is robust in the presence of a soma. Fig 7d illustrates that the number of dendrites that maximises the firing rate is greater for lower *ρ*_1_ and higher *μ*. We have discussed previously why the value of *n* that maximises firing increases with *μ* as the increase in mean from additional dendrites becomes more significant for the firing rate. Decreasing *ρ*_1_ increases the value of *n* that maximises firing because the relative increase in conductance by adding another dendrite is smaller when the fixed somatic conductance is larger.

## Discussion

This study demonstrated how the spatio-temporal fluctuation-driven firing of neurons with dendrites, soma and axon can be approximated using the upcrossing method of Rice [27]. Despite being reduced models of neuronal structures, they demonstrate considerable richness in behaviour beyond what point-like or compartmental models capture. For the one and two-dendrite models, the firing rate was shown to be independent of the electrotonic length constant; given that the length constant sets the range over which synaptic drive contributes to voltage fluctuations, this result is surprising. However, a dimensional argument extends this independence to any model in which semi-infinite neurites are joined at a point and share the same λ (any other properties without dimensions of length can be different in each neurite). The level-crossing approach provided a good approximation for the firing rate for these simple dendritic neuron models in the low-rate limit. Beyond this limit, simulations suggest that there is a universal functional form for the firing rate when parametrised by *σ_υ_* that is independent of both λ and the number of dendrites radiating from the nominal soma. This functional form, for coloured noise and in the white-noise limit, merits further mathematical analysis as it is distinct to that of the point-like integrate-and-fire model.

Extending the study to multiple dendrites, we showed that the firing rate depends non-monotonically on their number: adding more dendrites driven by fluctuating synaptic drive can, for a broad parameter range, decrease the fluctuation-driven firing rate. Dendritic structure has been previously shown to influence the firing rate for deterministic input [64, 65]. However, apart from the work of Tuckwell [30–32], analytical studies of stochastic drive in extended neuron models have largely focussed on a single dendrite with drive typically applied at a single point [36, 39] rather than distributed over the dendrite, or as a two-compartmental model [66]. This study demonstrates that in the low-rate regime, the upcrossing approximation allows for the analytical study of spatial models that need not be limited to a single dendrite nor with stochastic synaptic drive confined to a single point, but distributed as is the case in vivo.

Including axonal and somatic conductance loads demonstrated their significant effect on the firing rate - even relatively small axonal loads caused a marked reduction. Furthermore, the non-monotonic dependency of the firing rate on dendrite number was also shown to be affected by axonal radius and somatic size, demonstrating that the upcrossing method can be used to examine how structural differences in properties affect the firing rate of complex, composite, spatial neuron models.

The advantage of the level-crossing approach is it can be straightforwardly extended to include a great variety of additional biophysical properties affecting neuronal integration of spatio-temporal synaptic drive. An example of this for pyramidal neurons would be the inclusion of non-passive effects arising from voltage-gated currents such as *I_h_* [67]. The only requirement for the upcrossing approximation is the derivation of the voltage mean, variance and rate-of-change variance at the point that action potentials are triggered. For many scenarios, particularly in the high-conductance state, the spatio-temporal response can be approximated as quasi-linear, allowing the voltage moments to be calculated via Green’s functions using existing theoretical machinery, such as sum-over-trips on neurons [68–70]. The approach can also be extended to examine the dynamic firing-rate response to weakly modulated drive. This has already been done for point-neurons using the upcrossing method [29, 71, 72] and would only necessitate calculating the linear-response of voltage moments in the non-threshold case.

In summary, the extension of the upcrossing approach to spatially structured neuron models provides an analytical in-road for future studies of the firing properties of extended neuron models driven by spatio-temporal stochastic synaptic drive.

## Appendix

### Green’s functions

For notational simplicity we choose to measure distance in each neurite in terms of the length constant so *x_k_*/λ_*k*_ → *x_k_* and *y_k_*/λ_*k*_ → *y_k_*. In the Fourier domain, the Green’s function 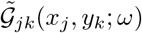 denotes the response at *x_j_* in neurite *j* to an input in neurite *k* at *y_k_*. The voltage due to an input in the Fourier domain 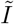 is given in terms of the double integral

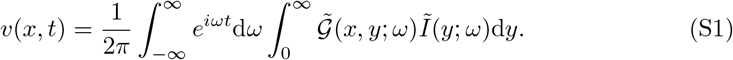

The Green’s function in the Fourier domain for Eq (18) satisfies the equation

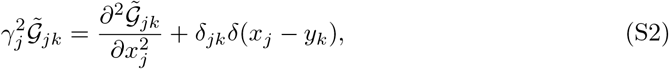

where *δ_jk_* is the Kronecker delta function. Eq (S2) has the general solution

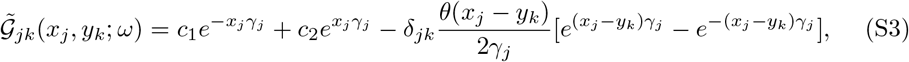

where the Heaviside step function *θ*(.) is only relevant if the input neurite and output neurite are the same. As the Green’s function inherits the boundary conditions of the system it describes, we apply a model’s boundary conditions to Eq (S3) to obtain the specific solution. Green’s functions for each of the cases studied in this paper are given later in this appendix. For an infinite dendrite, the Green’s function has the well-known form [73]

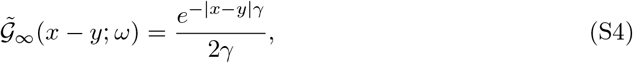

where 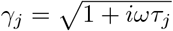. For multiple neurites and a soma, one can build more complex Green’s functions from a generalisation of Eq (S4) using the sum-over-trips formalism [68, 69]

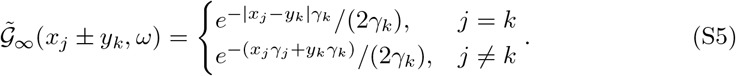

The only additional calculation that needs to be made is the segment factor 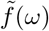. This quantity is the ratio of the admittance of the input neurite *Y_k_* (with *k=α* indicating the axon), to the total admittance of all neurites which radiate from the same node. For *n* dendrites radiating from a node with a soma this is

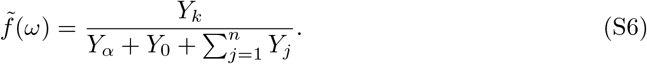

For each neurite, the admittance is given by

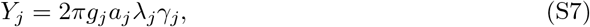

where *g_j_* is the total membrane conductance (including tonic synaptic conductance) while for a soma with membrane conductance *G*_0_, the admittance is

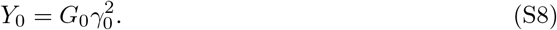

If there is only a single path from *x_j_* to *y_k_* and *j* ≠ *k*, then the Green’s function is given by Eq (S23)

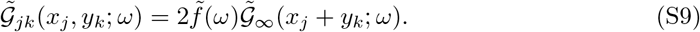

### Calculation of moments

The input *I* to each neurite has a deterministic and stochastic component, which in the Fourier domain are

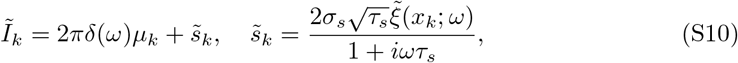

where we have again removed the units from distance. Since the system is linear, this means that the voltage will have a mean and fluctuating component

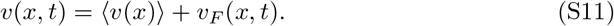

Substituting 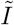 into Eq (S1) and taking the expectation, the mean in neurite *j* due to input in *k* is

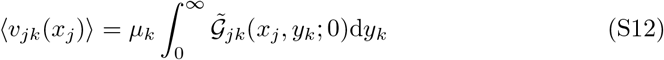

while the fluctuating component is

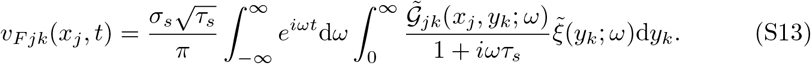

Thus the variance contribution from neurite *k* is obtained by squaring *υ_Fjk_* and taking the expectation, noting that 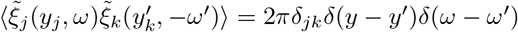,

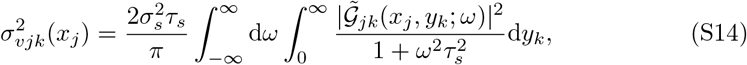

and similarly the variance of the voltage time derivative is found by multiplying the integrand of Eq (S14) by *ω*^2^

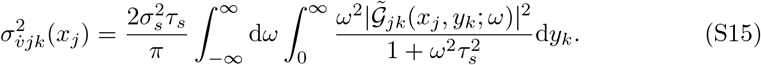

This approach is equivalent to that found in [74], where the integrand of Eqs (S14, S15) is proportional to the power spectral density of the voltage. With these integrals for the second moments, the definition 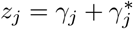 is useful for keeping the algebra compact.

For *n* dendrites with synaptic input, the response in the axon is simply the linear sum from each dendrite,

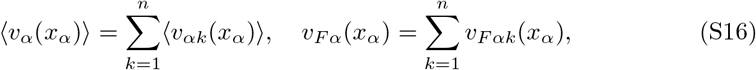

and since the stochastic drive between dendrites is uncorrelated, the second moment contributions from each dendrite also sum linearly,

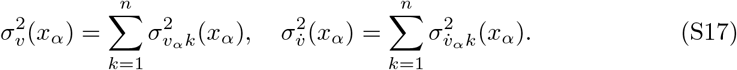

For dendrites with identical properties and drive, this means we can multiply Eqs (S14, S15) by *n* to obtain the total second moments.

In all the cases given, 〈*υ*〉 is easily analytically calculable. For the infinite dendrite 〈*υ*〉 = *μ*, while the resting potential in the axon for *n* dendrites is

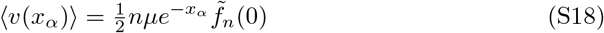

and the addition of a soma changes this to

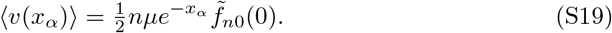

For many simple cases - such as the sealed dendrite, one-dendrite, and two-dendrite models - closed-form expressions for the second moments are attainable. For all cases with an axon and/or soma with different membrane properties to the dendrite, the *ω*-integral can be calculated numerically or approximated in a limit of interest. However, given the n in the denominator of 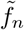 and 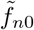, we expect from Eqs (S14, S15) that the second moments scale as ~ 1/*n* for large *n*.

### Derivation of Green’s functions

#### Closed dendrite

Given the zero-current boundary conditions at the ends *x* = [0, *l*]

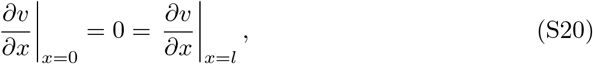

we can solve the Green’s function differential equation, Eq(S2), to obtain [73]

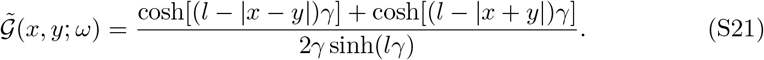

#### Dendrite and axon

Using the sum-over-trips method, 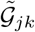 is given by the sum of infinite Green’s functions of each path which traces back from output position *x_j_* to input position *y_k_*. If a given path has length *l*_trip_, then we represent this sum as [68, 69]

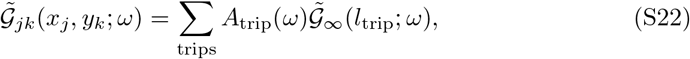

where *A*_trip_ is the trip coefficient that depends on the intersections between cables that a trip must path through. Since the neurites we consider are semi-infinite, there is only a single trip for a path from the axon to the input dendrite (however, the sum-over-trips approach provides a method for straightforward generalisation to dendrites with closed ends). The only trip coefficient required is that for transmission through a node which is given by 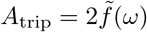 [69]. Therefore, the Green’s function from the dendrite *k*=1 to the axon, *j* = *α*, is given by

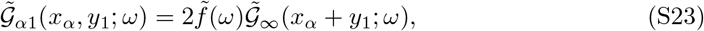

which upon substitution of 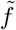 and 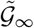 yields

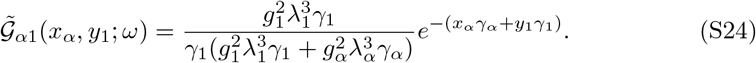

#### Multiple dendrites and axon

When there are *n* dendrites, the segment factor becomes

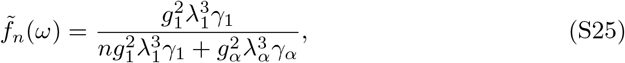

and hence the Green’s function for the axonal response is

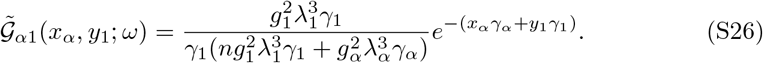

Since all dendrites have the same properties for this model, we can then claim that 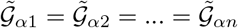.

#### Dendrites, soma and axon

For an electrically significant soma, the segment factor is now

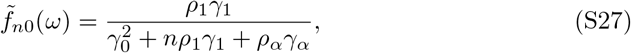

hence the Green’s function is

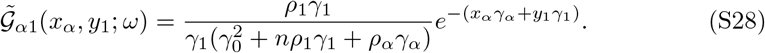

### Derivation of second moments

With an additive input in the Fourier domain 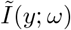, the voltage in the Fourier domain is given by

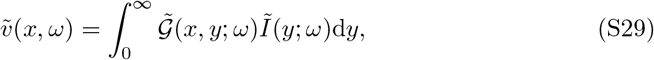

and after taking the inverse Fourier transform we obtain Eq (S1). While there are other methods for obtaining second moments that may be more convenient for the models which provide closed-form solutions (such as a Green’s functions in time [30] or Fourier series decomposition [30–32]) the method we present here extends most easily to arbitrary neuronal structures. For clarity of explanation, we derive the two-dendrite model first.

#### Two-dendrite model

For the two-dendrite model 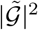 is given by

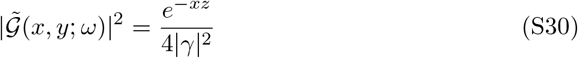

which we can readily integrate with respect to *y* after substituting into Eq (S14) to receive

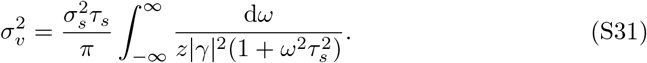

Using the substitution 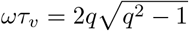 and splitting into partial fractions this integral becomes

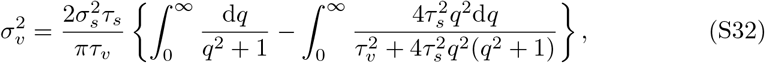

which can be resolved to give Eq (24).

#### One-dendrite model

Defining *iu* = *γ − γ*\* we find

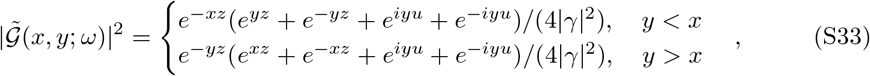

which integrates with respect to *y*, giving

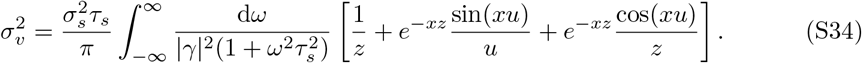

We can see that at *x* = 0 and as *x* → ∞ the variance is double and equal to the two-dendrite variance, respectively. For general *x* we can change the integration variable in a similar manner to the two-dendrite model to obtain the desired result.

#### Closed dendrite

With the closed dendrite, 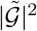 is more lengthy

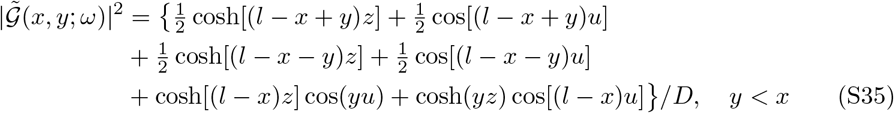

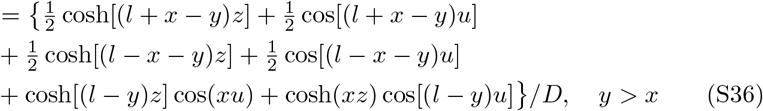

where *D* = 2|*γ*|^2^ [cosh(*lz*) − sin(*lu*)]; however, we can see that all the functions involved will integrate with respect to *y*.

#### Dendrite and axon

For the dendrite and axon, we will leave 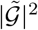 in terms of the segment factor to show how this approach extends to multiple dendrites and the addition of a soma

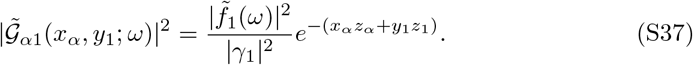

Integrating with respect to *y* gives,

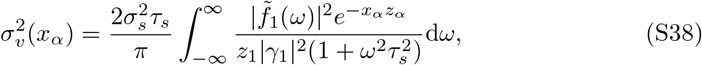

which generalises to *n* dendrites or the addition of a soma by replacing 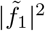 with 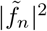 or 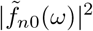 respectively.

### Calculation of axonal parameters

Assuming that the following parameters have the same value in the dendrite and axon: *E_L_, g_L_* and *c*_m_, we can express the axonal parameters in terms of the dendritic ones. Since there is no synaptic drive in the axon, *g_α_ = g_L_*, while in the dendrite *g*_1_ = *g_L_* + 〈*g_s_*〉. We denote the ratio between the membrane time constants as *ϵ*, which given constant *c*_m_ is

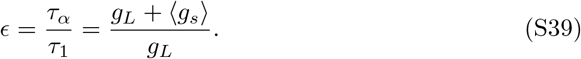

Recalling our definitions of *E* and *μ* in the dendrite as,

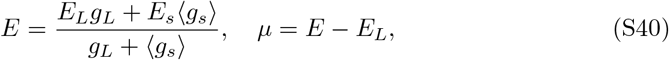

we can rearrange to find an expression for *ϵ* in terms of potentials alone,

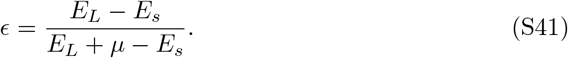

Hence we can calculate *τ_α_* in terms of *τ*_1_ given *μ, E_L_* and *E_s_*. For *E_L_* = −70mV, *E_s_* = 0mV and *μ* = 10mV this results in *ϵ* = 7/6.

When *a_α_* and *a*_1_ are fixed - as in Figs 5, 6c, and 7b - we can calculated λ_*α*_ in terms of a given λ_1_. Recalling the definition of the length constant from Eq (7) and making the reasonable assumption that the axial resistivity *r*_a_ is the same in the dendrite and axon, we have

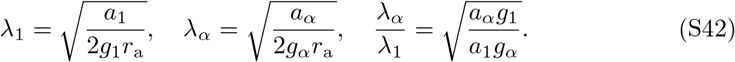

Taking *g_α_* = *g_L_* again and our earlier definition of *ϵ* in Eq (S39), we can write this as

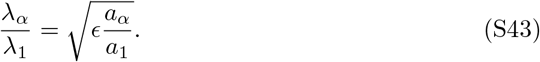

Finally, for the electrically significant soma in Fig 7 we give *ρ*_1_ but not *ρ_α_*, noting that it can be calculated given λ_1_ and λ_*α*_ or *a*_1_ and *a_α_*. Recalling our definitions of *ρ* and *ϵ* we find

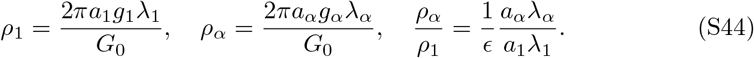

Hence when λ_1_ and λ_*α*_ are fixed as in Fig 7a,c,d we have

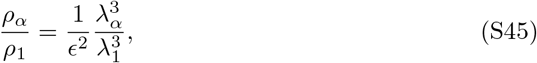

while when *a*_1_ and *a_α_* are fixed as in Fig 7b

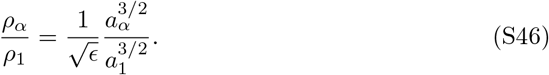

**Table S1.** Parameters and their default values used in the figures present in the main text. Since many of the parameters are inter-dependent, where a value is not given, a formula for how it is derived from the other parameters is given instead.

| Parameter | Units | Values used/Formula |
| --- | --- | --- |
| $E_L$ | mV | -70 |
| $E_s$ | mV | 0 |
| $\mu$ | mV | 4-12 |
| $\epsilon$ | - | $(E_L - E_s)/(E_L + \mu - E_s)$ |
| $\tau_1$ | ms | 10 |
| $\tau_\alpha$ | ms | $\epsilon\tau_1$ |
| $\tau_s$ | ms | 5 |
| $\lambda_1$ | $\mu\text{m}$ | 200 |
| $\lambda_\alpha$ | $\mu\text{m}$ | 100, 150, 200 |
| $a_\alpha/a_1$ | - | $g_\alpha\lambda_\alpha^2/(g_1\lambda_1^2)$ |
| $\rho_1$ | - | 1-16 |
| $\rho_\alpha$ | - | $\rho_1\lambda_\alpha^3/(\lambda_1^3\epsilon^2)$ |
| $\sigma_s$ | mV | 1-3 |
| $v_{\text{th}}$ | mV | 10 |
| $v_{\text{re}}$ | mV | 10 |
| $x_{\text{th}}$ | $\mu\text{m}$ | 0, 30 |

